# Family dynamics reveal that female house mice preferentially breed in their maternal community

**DOI:** 10.1101/2021.01.29.428775

**Authors:** Julian C. Evans, Anna K. Lindholm, Barbara König

## Abstract

Whether females breed in their natal group is an important factor in the evolution of extended families in animal sociality. Breeding in natal groups comes with clear costs and benefits, depending on size of the group and presence of older relatives, including mothers. Studying individual decisions about whether to stay or leave can provide insight into the mechanisms and trade-offs governing the formation and structure of family groups. We investigated the family dynamics of a large population of free-ranging commensal house mice. Using dynamic community detection on long term datasets, we determined which females first bred in their natal group. We then looked at how this influenced breeding success. We found most females (77%) exhibited strong philopatry, breeding in their natal groups. Whether a female bred elsewhere was only predictable when natal groups were extremely small and related or large and unrelated. Despite this preference, breeding elsewhere made no difference in how quickly and successfully a female bred. However, presence of their mother did lead females to breed sooner when born during high breeding activity, when competition over reproduction is high. Based on these results, potential loss of fitness does not seem to be the main driver of philopatry in female house mice. The effect of the presence of mothers may indicate retaining prior social connections is an important benefit of breeding in the natal group. Mothers providing benefits also suggests lack of conflict between generations, which is likely an important attribute in the development of extended family groups.

**Lay summary:** Whether animals breed in the group they are born in influences how they form extended family groups. Whether females stay will depend on properties such as presence of older relatives, including mothers. Using long-term wild mouse data, we track groups and which group females bred in. Most stayed, but leaving didn’t reduce breeding success. Presence of mother, who generally stayed, did lead to earlier breeding. This might be a key advantage to remaining to breed.

## Introduction

The formation of family groups is considered an important step in the evolution of animal sociality (Emlen, 1995; Kramer and Meunier, 2019). Family groups, defined as “an association of one or both caring parent(s) with their offspring after hatching or birth that arose and/or is currently maintained to enhance the fitness of the constituent individuals” (Kramer and Meunier, 2019), can be found in a large number of vertebrate and invertebrate species. The development and behaviour of a species’ juvenile offspring is a key part of the formation of such groups. In many cases, upon reaching nutritional autonomy or maturity offspring will disperse and seek independent opportunities to reproduce, rarely interacting further with their parents. In some species, however, offspring stay in contact with parents even after reaching maturity (Clutton-Brock and Lukas, 2012; Emlen, 1997). Therefore, while a “nuclear family” consists solely of parents and offspring from a single reproduction event, an “extended family” consists of individuals of varying degrees of relatedness, belonging to several generations (Kramer and Meunier, 2019). Within extended families, dominance asymmetries often occur between generations (Emlen, 1997; le Roux et al., 2011; Nelson-Flower et al., 2018), with older dominants controlling reproduction (Clutton-Brock et al., 2010) either physiologically (Massey and Vandenbergh, 1980; Young, 2009) or via despotic behaviours, such as infanticide (Clutton-Brock et al., 2001; Johnstone and Cant, 2010; Thompson et al., 2016). The structure of family groups will therefore affect reproductive strategies (Schradin and Lindholm, 2011), pervasiveness of polygamy and extra-pair copulations (Baglione et al., 2002; Tarvin et al., 2005; Wittenberger, 1980) or the evolution of associated adaptations in both sexes (Clutton-Brock, 2016). As such, whether offspring breed in their family group can influence population demography and dynamics, generating both negative and positive correlations between group size, survival and breeding success (Clutton-Brock and Lukas, 2012; Isbell, 2004; Kramer and Meunier, 2019).

Choosing to breed in a natal group comes with the costs of direct competition with relatives for resources (Clutton-Brock, 2017; West et al., 2002), as well as an increased risk of inbreeding depending on both sexes’ reproductive strategies. The increased competition with kin as offspring reach sexual maturity will increase the likelihood of offspring dispersal in some species (Gandon, 1999; Moore et al., 2006; Sorato et al., 2016). However, there are also many examples of offspring remaining in their natal group to breed. In some cases this may be due to ecological constraints preventing easy dispersal, or making the intrinsic benefits of grouping behaviour invaluable (Emlen and Vehrencamp, 1983; Hatchwell and Komdeur, 2000; Kokko et al., 2001; Komdeur, 2006). On the other hand the high level of relatedness within a family group can enable cooperative behaviours such as group foraging, group defence or alloparenting (Cockburn, 1998; Kaiser et al., 2018; Khodaei and Long, 2019; Ruch et al., 2009; Sherman, 1981). Alloparenting behaviours in family groups can range from communal care of offspring by multiple females (Ferrari et al., 2019; Vehrencamp, 1978) to non-breeders acting as helpers to breeding individuals (Clutton-Brock et al., 2006; Kaiser et al., 2018; Koenig et al., 1998; Schubert et al., 2009). If ecological constraints allow dispersal however, a young subordinate might benefit more in the short term by choosing to leave its natal group, engaging in independent reproduction without risk of reprisals, or potentially gaining a better dominance rank in a new group (Emlen and Vehrencamp, 1983; Johnstone, 2000; Keller and Waller, 2002; Komdeur, 2006; Stiver et al., 2004).

The decision of individuals to breed in their natal group can therefore provide insight into the mechanisms and trade-offs governing the formation and structure of family groups. Studying these decisions can be challenging however, generally requiring stable social groups where individual animals can be monitored over long periods of time. Because of this, many studies of family groups have typically focused on larger, long lived mammals (Goldenberg and Wittemyer, 2018; Johnstone and Cant, 2010) or on birds (Covas and Griesser, 2007; Griesser et al., 2006). Studying these dynamics in small mammals such as rodents is more difficult and therefore rarer as groups are often subject to fluctuations in size and composition, caused by variations in fecundity, mortality and emigration. Nevertheless, these species can display a high degree of variation in their social systems (French, 1994; Schradin, 2013; Schradin et al., 2012). Here we investigate the extended family dynamics in a large population of free-ranging commensal house mice (*Mus musculus domesticus*) in a barn in Switzerland. We examine the factors affecting female decisions to breed in their natal group after reaching sexual maturity and how this decision affects their breeding success. Female house mice typically live in social groups with overlapping generations, demic structure and female philopatry (Baker, 1981; Gerlach, 1996; König and Lindholm, 2012; Lewontin and Dunn, 1960). In our study population, groups are relatively stable over time and have proven to be resilient even after a major disturbance (Evans et al., 2020a; Liechti et al., 2020). While there is strong intrasexual female reproductive competition, several females usually breed simultaneously in the same group and even engage in communal offspring care (Ferrari et al., 2019). Female sociality has been suggested to enable this communal alloparenting (Weidt et al., 2014) and to protect altricial offspring from infanticide by non-group members (Auclair et al., 2014; Weidt et al., 2014). The study population is not food limited, meaning that competition for food with relatives is unlikely to be a reason to leave the natal group, though individuals might compete over mating opportunities (Coombes et al., 2018; Manser et al., 2020; Stockley and Bro-Jørgensen, 2011) or spaces to rear litters (Harrison et al., 2018). Similarly, as the barn is largely free from predators, meaning predation risk is unlikely to be a major cost of leaving the natal group. Therefore, loss of existing social connections and the need to successfully establish new ones to integrate with a new group is likely to be one of the primary costs of inter-group dispersal.

Using social network analysis allows us to monitor social groups and females’ relationships with them over time and multiple generations. We investigate females’ likelihood to breed in their natal group and the factors that might cause them to breed in another group. We then examine if this decision and the presence of their mother (as a close relative of the older generation in their chosen group) affects how quickly females successfully breed and their reproductive success. We predict females will preferentially breed in their natal group. However, since intragroup competition is expected to increase with number of adult females in the natal group, larger natal groups will result in a higher likelihood for females to breed elsewhere. Similarly, we also predict that high average relatedness to the other females in the natal group will increase a female’s probability of leaving to breed elsewhere, so as to avoid competing with the mother or other older relatives for mating or breeding opportunities. This potential for competition with older relatives in natal groups also leads us to predict that females who remain in their natal group will breed later, particularly if their mother is still present, due to inter-generation conflict. However, as females remaining in their natal group may benefit from a stable social position and established social relationships, we predict that those who remain in their natal group will have higher reproductive success than those who breed in another group.

## Methods

### Study population

The study population of commensal house mice was founded in 2002, in a 72m^2^ barn near Zurich, Switzerland. For a detailed description of the building and set-up see König and Lindholm (2012); König et al. (2015), see supplementary figure 1 for map. The environment found in this former agricultural building is typical of that experienced by house mice living in stables and barns throughout Europe. Food, water and nest building material are provided ad libitum. Temperature is continuously recorded inside the building on an hourly basis. Mice can exit and enter the barn freely, and inside the building they have access to 40 artificial nest boxes. The population is closely monitored, with all nest boxes regularly checked for new litters. Shortly before pups become mobile (13 days after birth ± 1 day), pups have genetic samples taken via an ear punch, which are used for parentage analyses. Although actual weaning begins at 17 days, the number of offspring raised to day 13 has been shown to highly correlate with weaning success (see Gerber et al., 2021). Approximately every 7 weeks, an attempt is made to capture all mice in the barn. As well as having a second genetic sample taken, adults weighing >17.5g are fitted with a passive integrated transponder (PIT) tag. These tags can be detected by the radio-frequency identification (RFID) antennae at the entrances to 40 artificial nest boxes. These antennae detect whether a mouse with a PIT tag is entering or leaving a nest box, from which it can be established which tagged individuals share nest boxes and for how long. Data were collected under the permits ZH 210/2003, ZH 215/2006, ZH 51/2010, ZH 56/2013 and ZH 91/2016 from the Cantonal Veterinary Office Zürich, Switzerland.

At tagging females are generally sexually mature but have not yet begun to breed (Carlitz et al., 2019; Ferrari et al., 2019; König and Lindholm, 2012). As juvenile female mice have been shown to have restricted ranges within their parental territory and avoid unfamiliar odour cues (Hurst and Nevison, 1994), we therefore find it unlikely that focal females left and then returned to their natal group to begin breeding.

Adult genetic samples are linked to pup genetic samples using markers at 25 polymorphic microsatellite loci (see Auclair et al., 2014; Evans et al., 2020b; Ferrari et al., 2019 for details). As pups can be aged with a high degree of accuracy (Ferrari et al., 2019; Gerber et al., 2021), this allows adults who were initially found as pups to be assigned reliable birth dates. The same genetic markers are also used to calculate the Wang coefficient (Wang, 2002) of pairwise relatedness among mice using the R package relatedness (Pew et al., 2018), an R implementation of the software COANCESTRY (Wang, 2011). Within this population, this measure has been found to correlate highly with Hamilton’s degree of relatedness *r*, with a Hamilton’s *r* of among full siblings *r*=0.5 corresponding to a Wang relatedness estimate of *r*=0.53 ± 0.02 (Harrison et al., 2018). Maternity is assigned using CERVUS 3.0 (Kalinowski et al., 2007), following Auclair et al. (2014). Assigned parentage was considered reliable at a 95% level of confidence, and if there was zero or only one mismatched allele between offspring and putative mother.

While litters that fail entirely can be discovered when a dead pup is found and successfully assigned to a parent, we think it highly unlikely we manage to detect all such instances. We therefore excluded the few of these failed litters we found from our analysis, so as to avoid bias. This means that, for the purpose of this study, a female was considered breeding when she raised the first litter (at least one pup) to the age of sampling at day 13 (range 12-14 days), and the number of offspring raised to day 13 was used as a proxy of breeding success.

### Network construction

For this study, we selected breeding females from 2007 to 2019 where the breeding female’s mother had a PIT tag when the female was born and the female had a PIT tag during their first recorded breeding event. For each focal female that met these criteria, we began by taking antennae data in a 31-day time period centred on the birth of the female (15 days on either side; female birth window). We then continued obtaining antennae data in 31-day time windows, until reaching the time window which contained the birthdate of the focal female’s first litter raised until day 13 (female breed window). Finally, we also took two time windows prior to the first (female birth) time window to better establish dynamic community structure at the female birth time window. Therefore, for each focal female, there was a minimum of four time windows (two pre-female birth, female birth, female breed), with a varying number of transition time windows between the birth and breeding time windows (transition). For an illustration of these time windows, see Figure 1.

**Figure 1:**
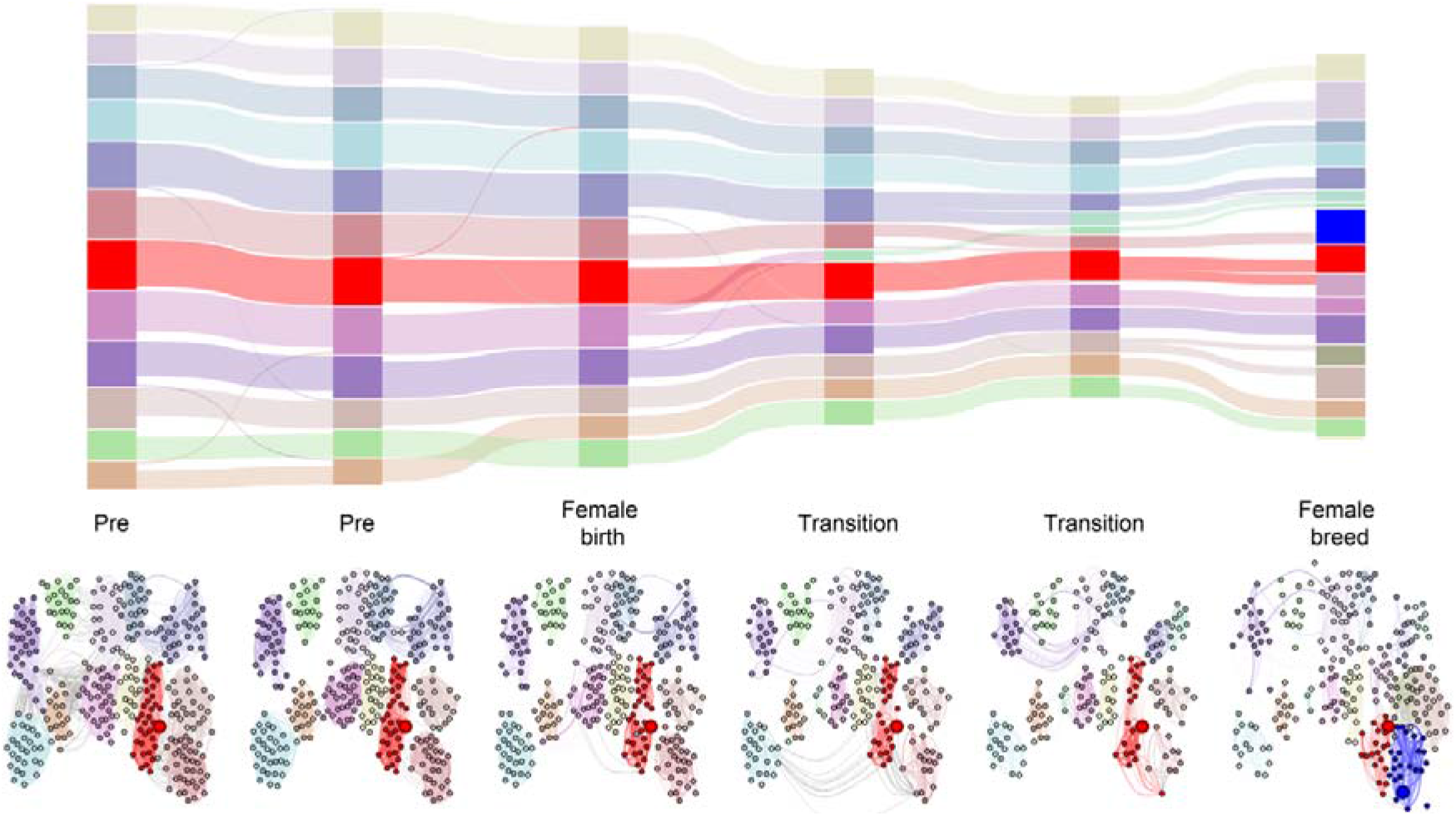
Example of networks and associated dynamic communities for a single focal female, who moved to a different group to breed (raise the first litter to day 13). The alluvial plot shows the detected dynamic communities per time window and how they relate to each other. The red community is the female’s natal group, while blue is the group in which they bred. Node colour in the network diagram corresponds to the alluvial plot, as do within-group edges. The large node with a red outline represents the focal female’s mother. The focal female is the large node with a blue outline. Between group edges are shown in grey. Edge width represents association strength. Node position is roughly based on that individual’s average location within the barn (see SI figure 1 for barn map and SI figure 2 for example of a similar plot where a focal female bred in their natal group).

For each time window, a social network was constructed. The antennae data within each of these time windows was used to construct a network consisting of every tagged mouse in the barn active for at least 20 days out of the 31 day period. This criterion was to ensure the accurate assessment of social associations and obtain better results when carrying out community detection. Edge weight was based on the time a pair of individuals spent sharing nest boxes, divided by the time individuals spent sharing nestboxes and the time each individual spent in nestboxes without the other (simple ratio index, where 0 indicates that individuals spent no time together and 1 indicates that individuals spent all their time in nest boxes together, Cairns and Schwager, 1987). For each network, community detection was carried out using the clustering algorithm developed by Blondel et al. (2008). This starts by assigning each individual their own group, and then sequentially moves allocated individuals between groups so as to achieve maximum modularity (proportion of connections within groups compared to between groups). All networks were constructed in R (R Development Core Team, 2020) using the igraph package (Csardi and Nepusz, 2006).

For each female we also calculated the average position in the barn in the female breed window, based on the coordinates of nest boxes weighted by how much time a female spent in each box in that time window. This was also calculated for the female’s mother during the female birth window. The distance between these average positions was used to estimate the change in home range of a focal female after birth.

### Dynamic community detection

We linked the communities detected in each time window into dynamic communities (i.e. communities that persist over time) using the MajorTrack python library (Liechti, 2020). This method (Liechti and Bonhoeffer, 2019) links detected communities between time windows using reciprocal majority identification. We used a two time window history parameter, meaning that for each time window the algorithm considered the community structure of the networks two time windows prior to that window. Therefore, for each female, we generated a dynamic community structure from the female birth window to the female breed window, with the two pre-female birth windows being used when carrying out dynamic community detection in the female birth window. Using this dynamic community structure, we identified a focal female’s natal group based on which dynamic community the female’s mother was part of when she gave birth to the female (female birth window). We then identified which dynamic community the female was part of when she gave birth to their first litter raised to day 13 (female breed window). Additionally, if the mother was still alive and present in the network during this time, we also identified which dynamic community she was a member of.

From the assigned dynamic communities, we determined if a female had moved to a different community to the one they were born in. We could also determine if they were in the same dynamic community as their mother (if the mother was still present) or the same community their mother had moved to subsequent to their birth. Finally, we also determined whether the dynamic community in which the focal female was born was still present when she bred. As the dynamic community detection algorithm can potentially assign a new dynamic community ID when groups undergo fission or fusion (the larger community retains its ID while the smaller is assigned a new ID), we were careful to consider the origin of that community when defining whether a female had moved to a new dynamic community. If the community the focal female was detected in emerged as the result of a fission or fusion event involving the natal group, this event had to have taken place before the transition window immediately prior to the female breed window, to be classified as a move to a new community. This was to focus on cases where the focal female had made an active decision to move to a new community, rather than cases where they were assigned a different dynamic community by the algorithm due to their original dynamic community splitting or merging with another.

### Network permutations and comparisons

In order to carry out hypothesis testing we used four different types of node label permutations, on all networks associated with each focal female (Farine, 2017). The first type permuted all of a focal female’s time window networks entirely randomly, with any individual able to swap with any other individual in each of that focal female’s time windows. The second permutation type enforced consistency in swaps over time windows, meaning that the same two individuals would be swapped in each of a focal female’s time window networks. The third permutation type limited swaps either within dynamic communities or between connected dynamic communities (linked via movement of individuals between the dynamic communities, or by merge or split events). The fourth type restricted swaps in the same way while also enforcing consistency of swaps over each of a focal female’s time window networks. For all four permutation types, we carried out a total of 10,000 node swaps for each of a focal female’s networks, in 100 sets of 100 swaps.

For each permutation type, we compared the proportions of focal females detected as breeding in their group and/or with their mother to the same proportions calculated from the 100 random datasets. P values were based on the number of proportions that were greater than the true proportion, divided by the total number of datasets in the comparison. For p values less than 0.05, the real proportion was deemed to be significantly different from that expected due to more random association patterns and movements between groups.

### Statistical analyses

All models were fitted in R using the BRMS package (Bürkner, 2017). As focal females’ mothers could appear in the dataset multiple times, the ID of the female’s mother was fitted as a random effect. The year containing either the focal female’s birth or breed time window was also fitted as a random effect, depending on the question being addressed. All models used the average temperature (during female birth or female breed windows) as a proxy for season. Previous studies of this population have shown that temperature has a strong correlation with breeding activity (Evans et al., 2020b; Ferrari et al., 2019; Gerber et al., 2021; König and Lindholm, 2012). All continuous variables were mean centred and rescaled so that 1 was equivalent to 1 standard deviation of the original variable.

#### What factors affect the decision to breed in natal group?

Whether a focal female bred in their natal group was fitted as a binary response variable using a Bernoulli distribution. Average temperature during the female breed window, the number of other adult females as well as the average relatedness between the focal and other adult females in the natal group during that time window, and all 2 way interactions between them were fitted as explanatory variables. Year of female breed window was fitted as a random effect.

#### Does remaining in natal group change time to breed successfully?

The presence of the mother and whether or not a focal female had remained in their natal group was combined into a single categorical explanatory variable with 4 levels, natal group with mother, natal group without mother, different group with mother and different group without mother. While we initially fitted the presence of a mother and breeding in natal group as separate terms with an interaction between them, focal females that fell into the category of different group with mother caused model convergence issues due to the small number of these cases relative to other categories. We addressed this by combining these terms into a single categorical variable and excluding focal females in a different group with mother from the analysis. We considered the presence of the mother a definite indicator of an older, more dominant relative in the group. The number of days elapsing between a focal female’s birth and the birth of their first litter raised to day 13 was fitted as a response variable, with mean temperature during female birth window, age of mother at female birth, and the categorical variable (mother present and whether the female bred in their natal group) as explanatory variables, along with all 2 way interactions. As initial data exploration suggested structured variance (primarily variance decreasing as temperature increased), all 3 explanatory variables were also fitted as variance effects. This allowed the variance estimated by the model to change depending on the value of each of these variables. Year of focal female birth and mother ID were fitted as random effects.

#### Does breeding in natal group influence the number of pups raised to day 13?

The number of pups in a focal female’s first litter raised to 13 days was fitted as a response variable with a Poisson distribution. The time taken in days to breed (date of birth of first litter raised to day 13), categorical variable of breeding in natal group and presence of mother, number of other adult females in the group during the female breed window, as well as average temperature, and all 2 way interactions between explanatory variables, were fitted as explanatory variables. Year of female breed was fitted as a random effect alongside mother ID.

## Results

From 2002 to 2019 a total of 726 focal females were suitable for inclusion in our analysis (see supplementary figure 4 for number of individuals per year), each with a mean of 10.11 ± 3.06 time windows. A further 28 individuals had sufficient network data, but no pups that survived till day 13 and were therefore not considered. Of the 726 individuals, 581 (77%) were still in their natal group when breeding (raising the first litter to day 13). This proportion was significantly greater than that found in all four different types of permutation (p<0.001 in all cases, Figure 2). Natal groups consisted of on average 17 adult females when a focal female was born. Focal females that bred in their natal group had a mean change in average position within the barn of 32.27 cm ± 24.89 SD, while those who bred in another group had a mean change in average position of 73.30 cm ± 58.40 SD. Among focal females whose mothers were still present in the barn when they bred (304, 42% of all females), 241 (79% of females whose mothers were still present, 33% of all females) bred in the same group as their mother. The vast majority of these were within the female’s natal group (215 individuals, 89% of all females whose mother was still present in the barn when they bred, 30% of all focal females). All these proportions were significantly greater than those produced by all four sets of permutations (p<0.001 in all cases, SI Figure). Mean time to give birth to the first litter raised to day 13 was 221.16 days ± 95.7 SD and the mean number of pups at day 13 in these litters was 2.67 ± 1.58.

**Figure 2:**
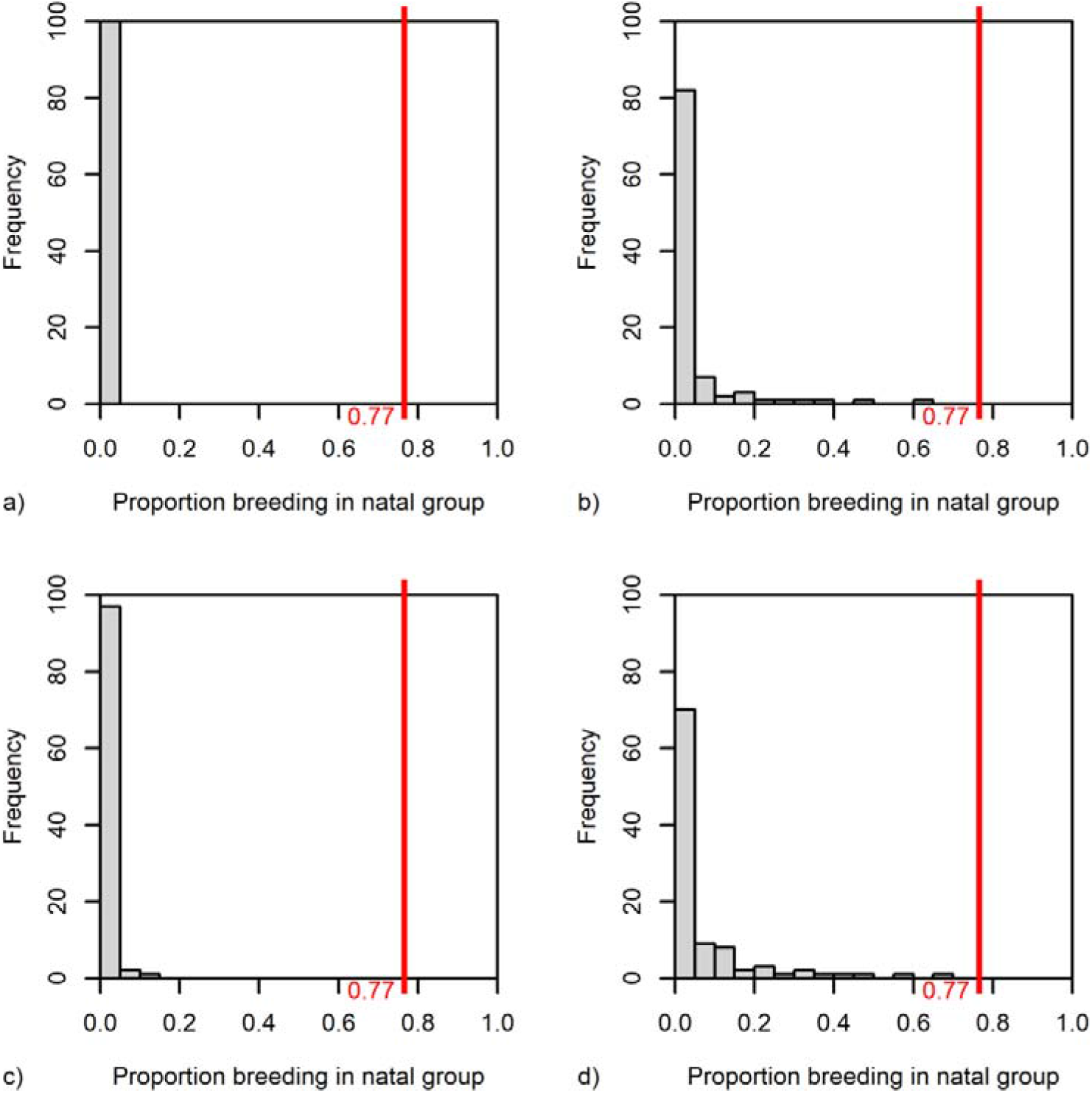
Frequency of the proportion of focal females breeding (raising the first litter to day 13) in their natal group in datasets permuted in four different ways: a) swapping individuals completely randomly in all time windows, b) swapping individuals randomly, but keeping swaps consistent between time windows, c) swapping individuals within their community and linked communities in all time windows, and d) swapping individuals within their community and linked communities while keeping swaps consistent between time windows. The proportion of focal females breeding in their natal group in the empirical dataset (77%) is indicated by the red line.

We also detected 237 cases where an individual was in the dataset both as a focal female and as another focal female’s mother. In 154 (65%) of these cases, the last mentioned focal female bred in the same natal group as their grandmother and mother (first focal female bred in their natal group where they gave birth to a second focal female who also bred in their natal group).

### What factors affect the decision to breed in a natal group?

The model examining what aspects of a focal female’s natal group at time of breeding might influence whether a female bred in their natal group found a strong interaction between the number of and average relatedness to other adult females in the natal group (Table 1, Figure 3). In groups with a large number of other adult females, the focal female was more likely to remain and breed the higher their average relatedness to the other adult females. In groups with a relatively small number of other adult females during a focal female’s birth window, the opposite effect was observed. The effect was negligible at an average number of other adult females in the natal group.

**Table 1:**
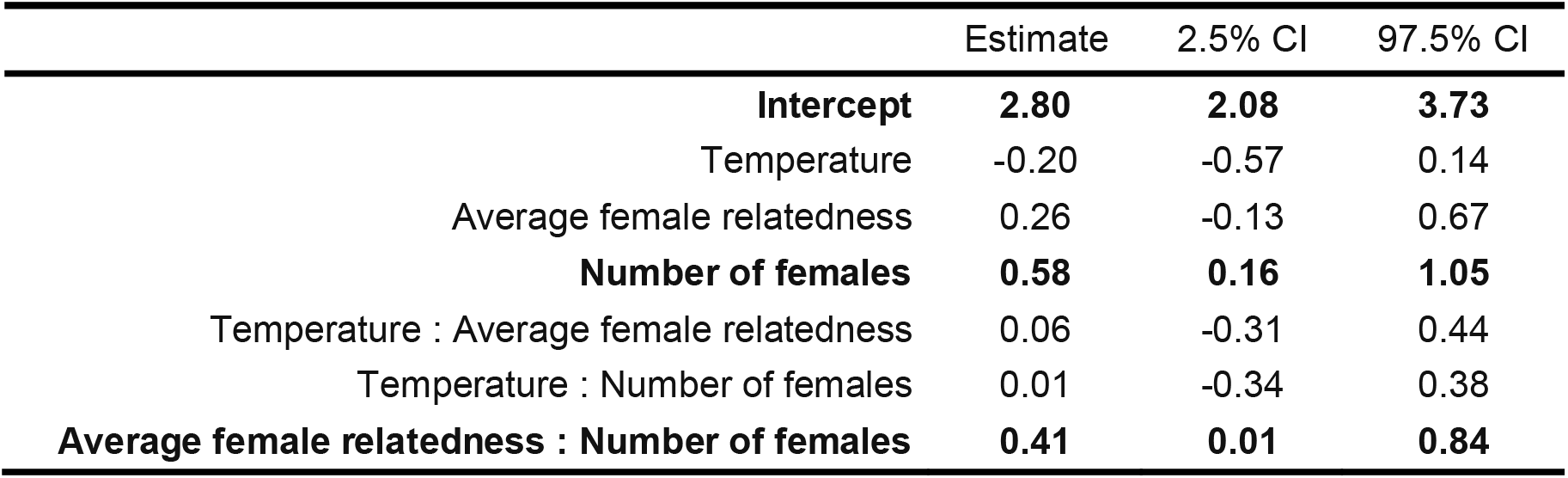
Model results for factors influencing a focal female’s probability of breeding (first raising a litter to day 13) in their natal group. Explanatory variables are the average temperature at breeding, average relatedness to other adult females and the number of other adult females in the natal group when breeding (female breed window; for time windows see Figure 1). Model was fitted with a Bernoulli distribution, ID of the female’s mother and year were fitted as random effects. All variables were mean centred and rescaled so that 1 = 1 SD of the unscaled variable. Bold text indicates a substantive effect (CIs do not cross 0).

**Figure 3:**
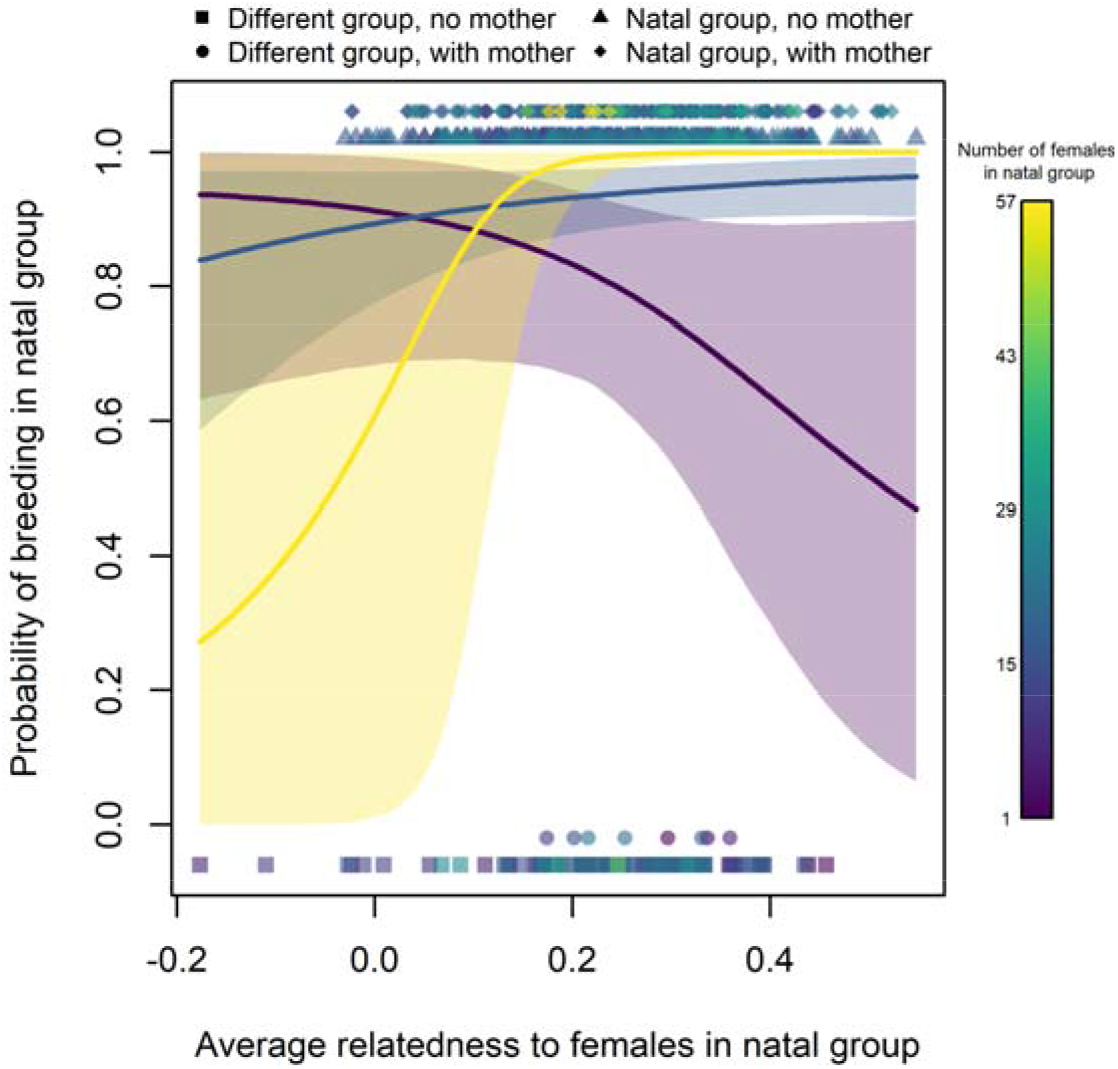
Probability of a focal female to breed (raise a litter to day 13) in their natal group (female breed window), compared to their average relatedness to all adult females (x axis) and number of other adult females in their natal group at breed window (colour of points and lines). Model estimates (Table 1) are plotted for the minimum (1), mean (16.79±7.12 SD) and maximum (57) number of other adult females in the natal group, with 95% confidence intervals. The four symbols indicate the presence of the mother in the group.

### Does breeding in the natal group affect the time to first raise a litter to 13 days?

Breeding activity in the study population generally correlated with temperature, with a first peak during spring (April to May, usually 2 to 20 °C), followed by high breeding activity in summer (June to August, usually 16 to 25 °C), and few litters born in winter (January to February, −2 to 8 °C). Focal females varied drastically in how long they took to give birth to their first litter raised to 13 days of age, depending on what time of year they were born and whether their mother was still present (mean 221.16 days ± 95.70 SD). Females born in a temperature range of 5-16 °C (within the range of typical spring temperatures) appeared to fall into two types. Those who bred early (mean 147.60 days ± 47.85 SD, values obtained from k-mean cluster detection of time to breed for focal females born in this temperature range) and those who took approximately a year longer (mean 348.90 days ± 60.33 SD, values obtained as above, Figure 4). Focal females born in warmer temperatures (summer, 17 °C and higher; mean 266.05 days ± 61.77 SD) did not appear to display such a split, indicated by negative temperature variance effect, indicating that variance in time to breed decreases as temperature increases (Table 2). Conversely, the few females born in colder temperatures (<5 °C, typical for winter months) tended to breed more quickly (mean 147.06 days ± 55.64 SD). The moderate number of focal females born in autumn (September to November, which had a similar temperature distribution to spring) followed similar patterns as those born in summer, usually taking longer to breed (210 days ± 55.06 SD). Focal females generally first bred earlier in spring if their mother was present, while those whose mother was not present were split between breeding early and late (Table 2, Figure 4). This was especially the case if the focal female’s mother was older when the focal female was born.

**Figure 4:**
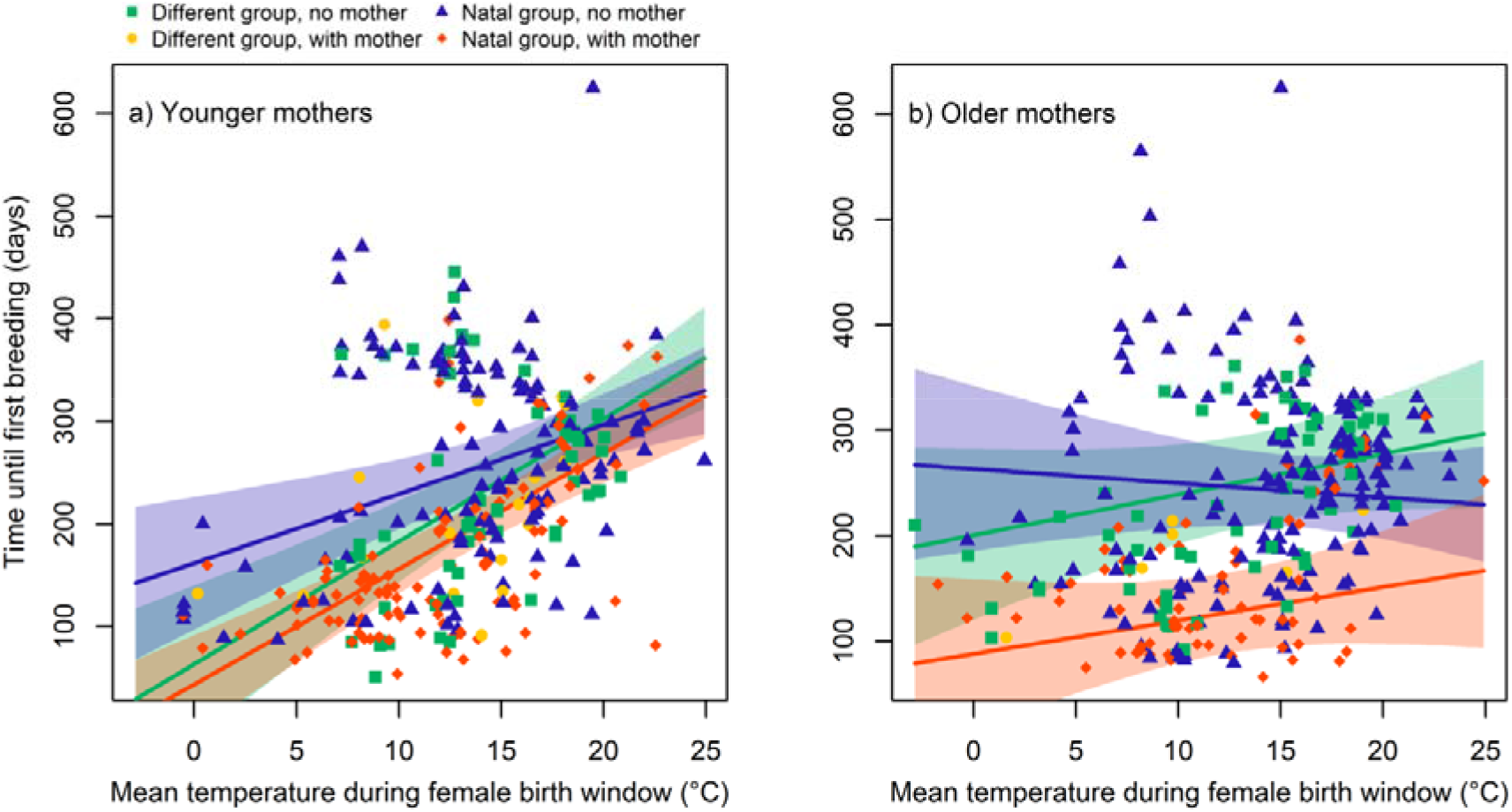
Influence of temperature at birth of a focal female (female birth window, x axis) and whether they bred in their natal group or not, along with presence of mother (colour and symbol), on time to first breed (raise the first litter to 13 days of age). a) shows data where age of the focal female’s mother was below average (293.9 days ± 115.6 SD), with model estimates at minimum age of a female’s mother (66 days), and b) shows data where the age of the focal female’s mother was above average, with model estimates at maximum age of a female’s mother (653 days). All model estimates presented with 95% confidence intervals. See supplementary figure 4 for a plot with all data in a single panel.

**Table 2:** Model results for the influence breeding in natal group and presence of mother have on how soon a focal female breeds (first raises a litter to day 13). Explanatory variables are the average temperature during the focal female’s birth (female birth window; for time windows see Figure 1), situation in which the female bred (in natal group with mother present, in natal group with mother absent, and in different group with mother absent) and the age of the female’s mother at their birth in days. These were also fitted as variance effects. ID of the focal female’s mother and year were fitted as random effects. Cases where females bred outside their natal group but with their mother present were excluded as the relatively small number of cases caused convergence issues. All variables were mean centred and rescaled so that 1 = 1 SD of the unscaled variable. Bold text indicates a substantive effect (CIs do not cross 0).

|  | Estimate | 2.5% CI | 97.5% CI |
| --- | --- | --- | --- |
| **Intercept** | **169.14** | **158.32** | **179.80** |
| **Different group, Mother absent** | **64.96** | **46.45** | **83.15** |
| **Natal group, Mother absent** | **80.08** | **64.16** | **95.74** |
| **Temperature** | **41.78** | **30.99** | **52.76** |
| **Mother age** | **-12.52** | **-23.66** | **-1.44** |
| Different group, Mother absent : Temperature | 3.40 | -16.71 | 22.97 |
| **Natal group, Mother absent : Temperature** | **-23.29** | **-39.83** | **-6.76** |
| **Different group, Mother absent : Mother age** | **18.19** | **0.71** | **35.87** |
| Natal group, Mother absent : Mother age | 11.32 | -4.31 | 26.97 |
| **Temperature : Mother age** | **-8.23** | **-15.06** | **-1.33** |

Variance effects
|  | Estimate | 2.5% CI | 97.5% CI |
| --- | --- | --- | --- |
| **Intercept** | **3.85** | **3.65** | **4.05** |
| **Different group, Mother absent** | **0.36** | **0.12** | **0.62** |
| **Natal group, Mother absent** | **0.56** | **0.35** | **0.78** |
| **Temperature** | **-0.18** | **-0.29** | **-0.08** |
| Mother age | -0.01 | -0.09 | 0.08 |

### Does breeding in natal group influence the number of pups raised to day 13?

No clear effect of breeding in natal group was found on the number of offspring in a focal female’s first litter raised to 13 days (Table 3). This was true regardless of mother presence, at all times of the year. However, those focal females breeding in a group containing a larger number of adult females raised fewer pups.

**Table 3:** Model results for the effect of breeding in natal group and presence of mother on the size of the first litter raised until day 13. Explanatory variables are the situation in which the focal female bred (in natal group with mother present, in natal group with mother absent, and in different group with mother absent), mean temperature at breeding (during female breed window; for time windows see Figure 1), number of other adult females in the group when breeding, and time taken (in days) to breed. Cases where focal females bred in another than their natal group but with their mother present were excluded as the relatively small number of cases caused convergence issues. Model was fitted with a Poisson distribution. ID of the focal female’s mother and year were fitted as random effects. All variables were mean centred and rescaled so that 1 = 1 SD of the unscaled variable. Bold text indicates a substantive effect (CIs do not cross 0).

|  | Estimate | 2.5% CI | 97.5% CI |
| --- | --- | --- | --- |
| **Intercept** | **1.01** | **0.88** | **1.14** |
| Different group, Mother absent | -0.03 | -0.17 | 0.12 |
| Natal group, Mother absent | 0.03 | -0.09 | 0.16 |
| **Number of females** | **-0.11** | **-0.21** | **-0.01** |
| Time to breed | 0.08 | -0.04 | 0.20 |
| Temperature | -0.02 | -0.11 | 0.08 |
| Different group, Mother absent : Number of females | 0.01 | -0.13 | 0.14 |
| Natal group, Mother absent : Number of females | 0.01 | -0.12 | 0.14 |
| Different group, Mother absent : Time to breed | -0.05 | -0.21 | 0.11 |
| Natal group, Mother absent : Time to breed | -0.04 | -0.17 | 0.09 |
| Different group, Mother absent : Temperature | 0.01 | -0.13 | 0.16 |
| Natal group, Mother absent : Temperature | 0.04 | -0.08 | 0.15 |
| Number of females : Time to breed | -0.01 | -0.07 | 0.04 |
| Number of females : Temperature | 0.02 | -0.03 | 0.07 |
| Time to breed : Temperature | 0.00 | -0.05 | 0.06 |

## Discussion

We used dynamic community analysis with a long term dataset of wild commensal house mice to examine to what extent females will choose to breed in their natal group and the effect this has on the onset and success of breeding. We found that the vast majority of females were philopatric, breeding in their natal group. Whether an individual would breed in another group was only predictable if their natal group consisted of a few highly related females, or many unrelated females. Contrary to our predictions, whether a female bred in their natal group had no effect on how quickly they bred, and how many offspring they raised until day 13 in their first litter. However, the presence of the mother consistently led to females breeding sooner if born during the first peak in breeding activity (during spring), particularly if the mother was older. Therefore, while breeding in natal groups seems preferable in female house mice, the natal group itself did not appear to confer any immediate reproductive benefit. Conversely, breeding in another group did not seem to confer any significant cost. Instead, the primary advantage of remaining in the natal group seems to be due to the benefit of remaining with the mother and the opportunity to breed sooner.

### Why leave the natal group to breed in another group?

As expected, we found that most females chose to breed in their natal group. Adult females seemed to tolerate the breeding of young females born in their group, and did not delay their onset of breeding. This suggests a low level of conflict between generations, unlike in species where younger individuals staying in their natal group may have to wait for an opportunity to breed (Hager and Johnstone, 2004). The main exceptions were in extremely small or large groups. Females were less likely to remain and breed in small groups of highly related females. This type of group may indicate a lack of group stability, making them undesirable to breed in. These small group sizes might also indicate on-going fission events, meaning that most individuals move to other groups, regardless of their breeding status. Outside of these extreme cases, avoiding competition with relatives seemed generally not to cause females to breed elsewhere.

Our results also indicate that resource competition within the natal group did not force dispersal, since focal females were only predicted to breed elsewhere if their natal groups consisted of a large number of unrelated adult females. There is negligible competition over food within the study population. Additionally, multiple paternity is common in female house mice (in our study population, almost 50% of the litters are sired by more than one father; Auclair et al., 2014; Manser et al., 2020), and breeding females regularly roam into neighbouring territories (Hurst, 1987, 1990). This makes it unlikely that competition for mating partners would cause individuals to breed outside their natal group. Similarly, while litters born outside nest boxes have a very low chance of success (Harrison et al., 2018), females rarely lack the option to access empty nest boxes (Ferrari et al., 2019; Harrison et al., 2018), and can potentially overcome this limiting factor by pooling their litters and engaging in indiscriminate communal nursing (Auclair et al., 2014; Ferrari et al., 2019; Weidt et al., 2014). In our study, 520 of our 726 focal females raised litters communally, despite the fact that communal nursing decreases the number of offspring weaned due to exploitative female nursing partners committing infanticide so as to improve parental care for their offspring (Ferrari et al., 2019; König, 1994a). This can lead to asymmetry of breeding success between communally nursing females (Ferrari et al., 2019). The costs of such exploitation will be lower if the females are related, due to indirect fitness benefits of raising a relative’s offspring offsetting the reduced fitness due to infanticide. Relatives therefore might be more likely to tolerate reproductive asymmetry, allowing for some degree of cooperation even in the presence of exploitation (Mathot and Giraldeau, 2010). This may make it desirable to stay and breed in a more related natal group, so as to avoid being exploited by non-relatives. Regardless of how evenly distributed the costs and benefits of communal nursing are between nursing partners (Ferrari et al., 2019), it could lead to a more egalitarian social system among female house mice than we might otherwise expect, unlike species where females are forced to breed outside their natal group. However, while communal nursing could certainly facilitate breeding in the natal group, it is generally thought that philopatry evolves before cooperative behaviour (Clutton-Brock and Lukas, 2012; Nelson-Flower et al., 2018). This suggests further benefits to breeding in the natal group, or potential costs of leaving.

### Why stay and breed in the natal group?

In some species, individuals staying and attempting to breed in their natal group may have a greater chance of eventually inheriting another family members’ dominance rank and associated benefits, such as reproductive rights or territories (Gaston, 1978; Nelson-Flower et al., 2018; Ragsdale, 1999; Stiver et al., 2004; Waser, 1988). Breeding in the natal group and perhaps initially engaging in communal nursing might be considered making the best of a bad job, while waiting to gain greater benefits in future (Bergmüller and Taborsky, 2005; Gaston, 1978). Remaining in a natal group to breed could also allow individuals to take advantage of established social associations. Stronger social associations are thought to lead to a greater probability of engaging in cooperative behaviours (Clutton-Brock, 2002; Van Horn et al., 2004) and to reduce levels of aggression or infanticide (Carazo et al., 2014; König, 1994a; McComb et al., 2001; Pravosudova et al., 2001). In house mice, females prefer to nurse communally with more familiar individuals who may be more likely to tolerate their presence (Ensminger and Meikle, 2005; Harrison et al., 2018; Hurst and Barnard, 1995; König, 1994b; Parmigiani, 1989; Rusu and Krackow, 2004). The value of social associations might further increase the potential costs of leaving the natal group and failing to integrate with a new group (Cameron et al., 2009; Komdeur, 2006). It has been suggested that in some species, individuals will spend time associating with nearby groups, so as to ease the eventual joining of that group and avoid spending an extended period of time with no group (Bergmüller et al., 2005; Stiver et al., 2004). In our study, females that bred outside their natal group mostly moved to groups which were relatively close spatially. This might indicate that female house mice choosing to breed outside their natal group will still prefer a group with which they have some familiarity. Due to the lack of predators in the barn, the cost of leaving the natal group is highly unlikely to be related to predation risk, though perceived predation risk in less familiar areas might still be high (Debeffe et al., 2013; Hulthén et al., 2015).

Given the resource abundance in the barn, lack of familiarity with areas outside of the territory of the natal group also seems unlikely to be a cost, though this might depend on individual preferences (Cooper et al., 2017; Debeffe et al., 2013; Dingemanse et al., 2003). Similarly, breeding outside the natal group also seemed to have no effect on the onset of breeding or the size of the first litter raised to day 13. This suggests low costs of breeding in another group. Therefore, difficulty of integrating with a new group and the potential loss of social connections may be the primary cost of leaving the natal group to breed elsewhere.

### The presence of mothers matters

Despite the clear preference for breeding in the natal group, we found this had no effect on reproductive measures. We did, however, find that females sharing a group with their mother bred more quickly during periods of high breeding activity. Given that in the vast majority of cases the mother remained in the natal group, this may be a strong reason for females to attempt to breed there. While we focus here on the female’s first breeding, a simple model showed that lifetime reproductive success tended to be higher the younger a female was when they first raised a litter to 13 days of age (SI Table 1). A caveat about this result is that our dataset does not capture litters that failed entirely. Females who took longer to raise their first litters until day 13 may have had undetected unsuccessful litters in the interim. While breeding more quickly in the presence of the mother seems like clear evidence that there is no conflict with the mother, the exact advantage a mother might confer is uncertain. Still, the presence of mothers may assist females with the creation of social bonds (Armitage et al., 2011; Berman et al., 1997). Alternatively, a female may be able to inherit and take advantage of their mother’s social bonds, possibly giving access to support provided by associates in social conflict situations (Ilany and Akcay, 2016; Stanton and Mann, 2012). Our model also indicated that a female’s mother being older would further increase how quickly a female raised their first litter to day 13. Older, more experienced mothers could provide better parental care when the female is younger, increasing their likelihood of successfully mating and breeding more quickly. Older mothers might additionally possess more established stable social associations. An older mother being present might also simply be an indicator of the group being safe and socially stable, increasing a female’s chances of successfully breeding more quickly (Cameron et al., 2009; Wey et al., 2013).

### Conclusions

Our study utilises a detailed long term dataset to examine whether breeding inside or outside the natal group will have fitness impacts. We find that most females display a high degree of philopatry, but breeding outside the natal group seems to have no impact on breeding. However, the presence of a mother can lead to females breeding more quickly when they are born at the beginning of the major breeding season in spring, under conditions of presumed high reproductive competition. This suggests low conflict between generations, possibly assisted by females being able to engage in communal nursing. Further study of the benefits that the presence of mother can provide, social or otherwise, will be necessary to determine if this is the main driver of breeding in natal groups in this species.

Distinguishing whether benefits or costs lead to the high rate of breeding in the natal group observed here would require more details on the behaviour of females immediately after weaning. Ideally, individuals would be tracked from an early age so that any movement between groups before breeding would be recorded. This would require tagging individuals at a younger age than in this study. Furthermore, tracking individuals from such an early age would provide information on how much a female’s social contacts will derive from their mother. Similarly, how important a social contact their mother is during the early establishment of social bonds could be established by looking at the likelihood of females and their mothers forming mutual contacts. Detailed analysis of how social bonds are affected when an individual joins a new group, dependant on their level of contact with that group in the past would determine the importance of established connections in the natal group. Finally, greater knowledge of the exact ecological constraints affecting whether individuals leave their natal group would also help better understand what is influencing this decision.

## Acknowledgements

We thank the numerous research assistants who assisted with data collection over the study period used here, specifically Gabi Stichel, Sally Steinert and Bruce Boatman. We also thank Jari Garbely for carrying out genetic analyses of thousands of samples and Damien Farine for helpful comments.

## Author contributions

JCE devised the study, processed and analysed the data and wrote the manuscript. BK initiated and maintains the long-term study, AKL is involved in long-term data collection, and maintains the genetic data. All authors discussed the results and contributed to the final manuscript.

