## supplementary for "Family dynamics reveal that female house mice preferentially breed in their maternal community"

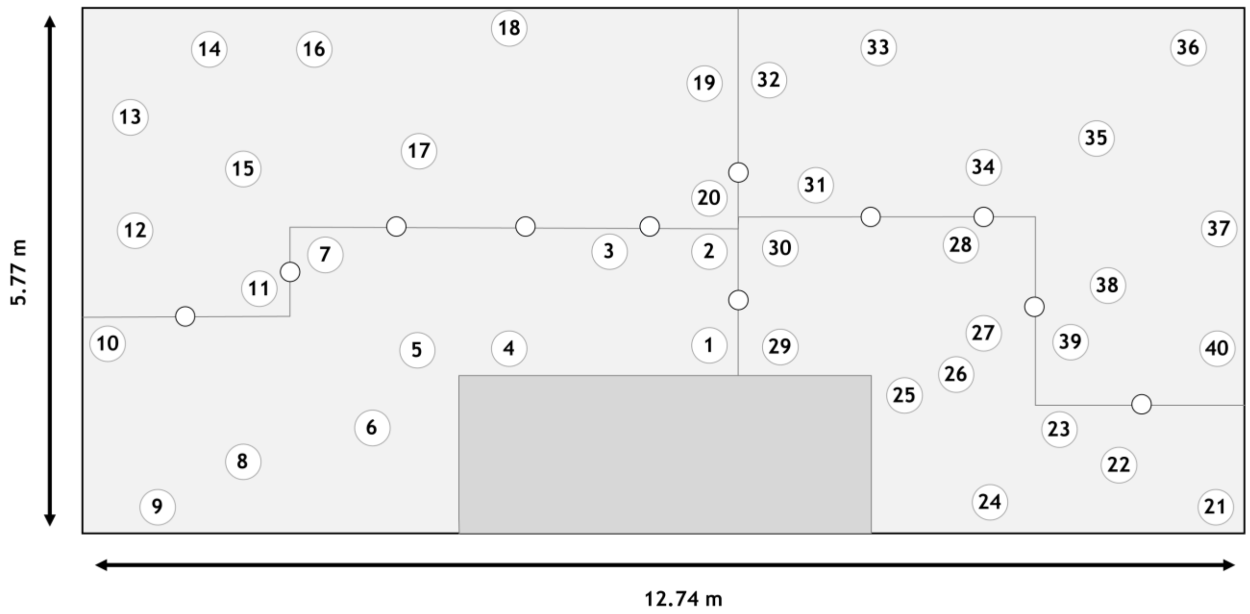


Supplementary figure 1: Map of the barn indicating location of RFID equipped nest boxes and barriers in separating the four sections, which mice can access through various holes in the barriers and by climbing over the barriers. Each section also includes 3 food bowls, 5 water bottles and a number of extra shelters and smaller barriers.

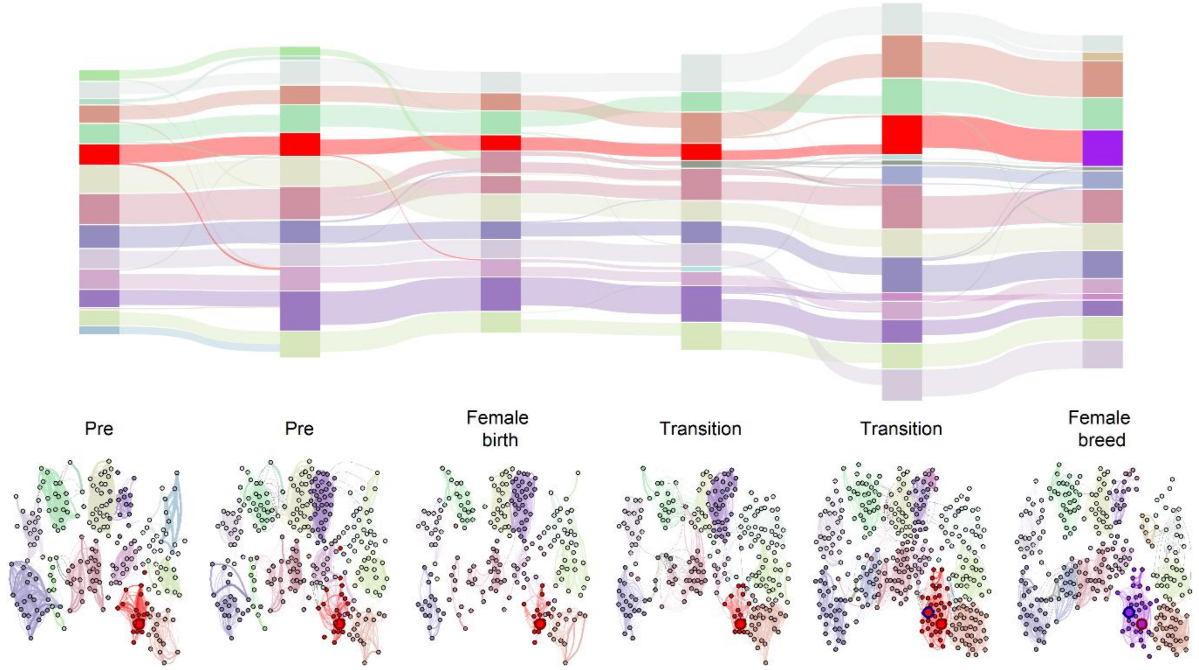


Supplementary figure 2: Example of networks and associated dynamic communities for a single focal female, who remained in their natal group to breed (raise the first litter to day 13). The alluvial plot shows the detected dynamic communities per time window and how they relate to each other. The red community is the female's natal group, while blue is the group in which they bred. Node colour in the network diagram corresponds to the alluvial plot, as do within-group edges. The large node with a red outline represents the focal female's mother. The focal female is the large node with a blue outline. Between group edges are shown in grey. Edge width represents association strength. Node position is roughly based on that individual's average location within the barn.

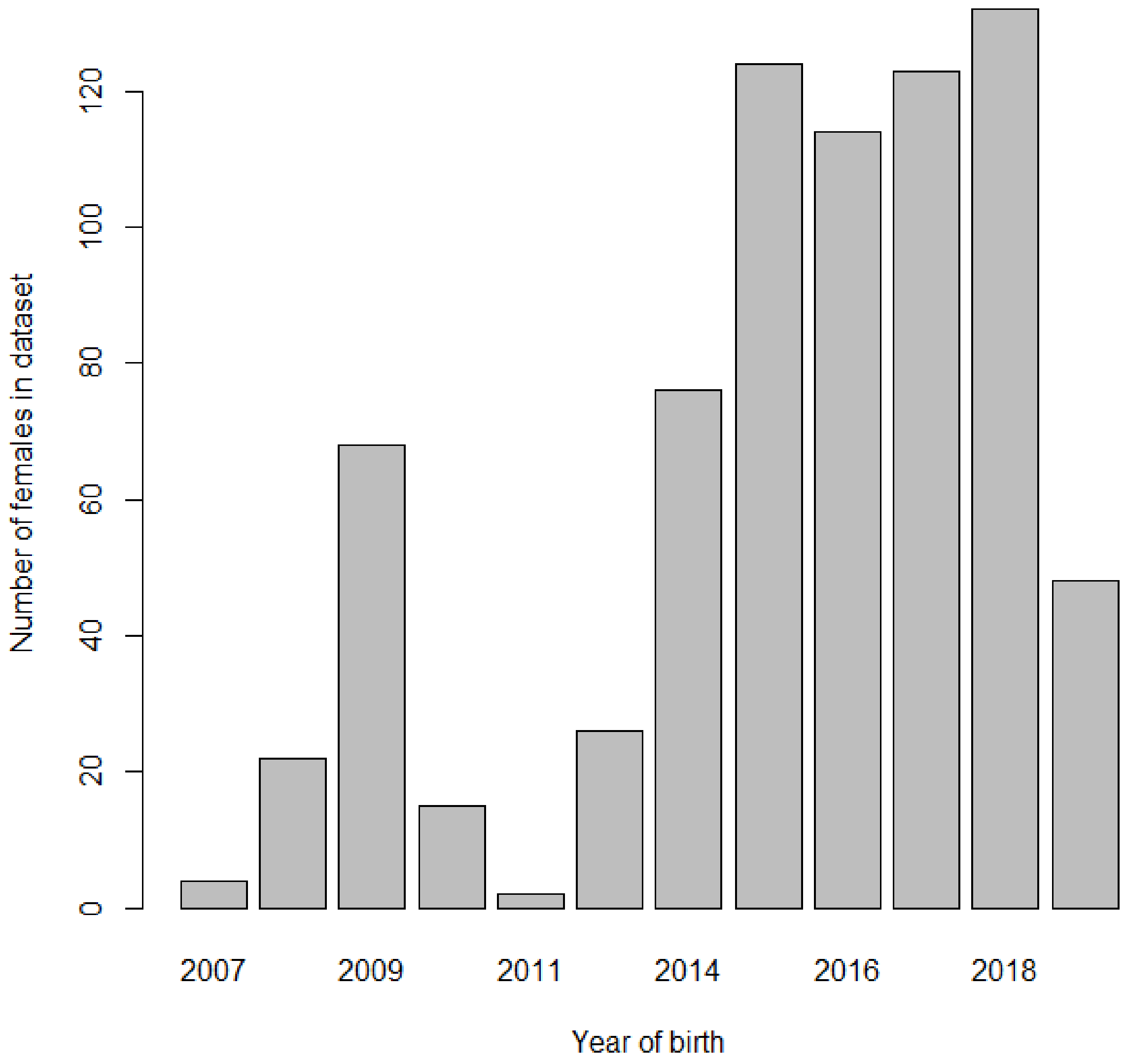


Supplementary figure 3: Number of females in dataset by the year they were born. From 2010 to 2011 the antennae system suffered an outage in which limited the quality of data. At the time of analysis, genetic samples had been processed through to mid 2019.

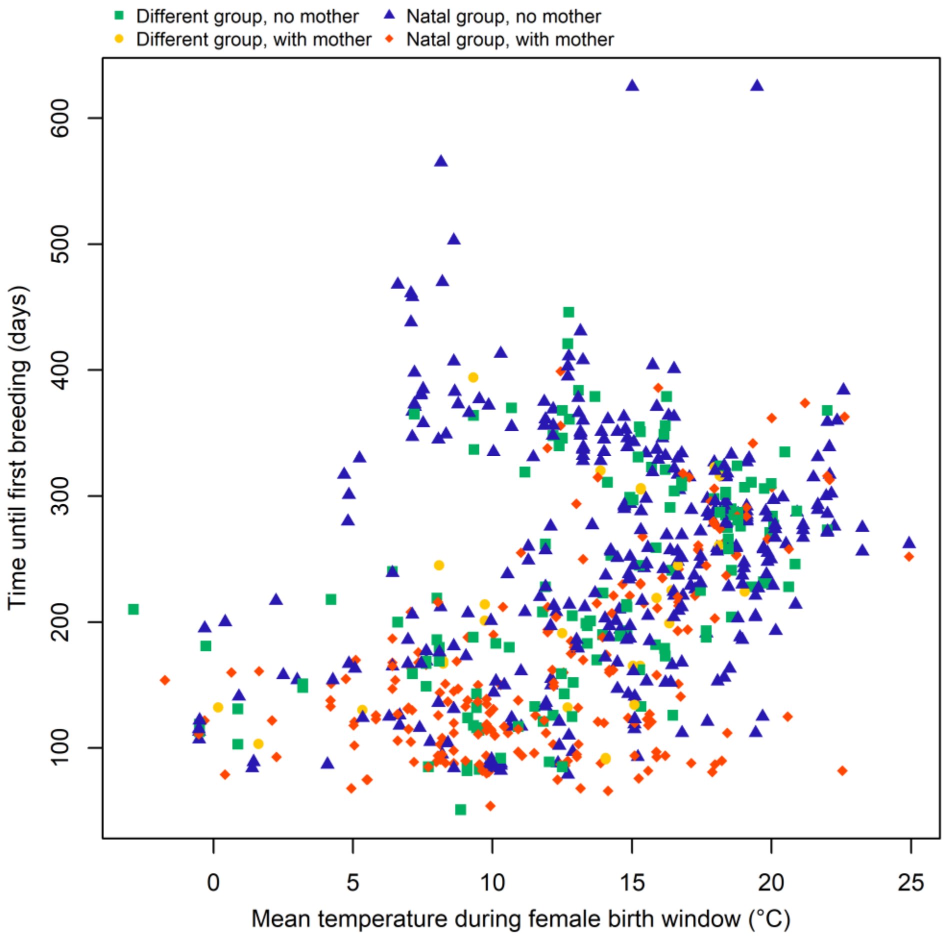


Supplementary figure 4: Temperature at birth of a focal female (female birth window, x axis) and whether they bred in their natal group or not, along with presence of mother (colour and symbol), plotted against time to first breed (raise the first litter to 13 days of age).

Supplementary table 1: Model results for the effect of first breeding time (days till first litter raised to day 13) on lifetime reproductive success. Model was fitted with a Poisson distribution. ID of the focal female's mother and year were fitted as random effects. All variables were mean centred and rescaled so that 1 = 1 SD of the unscaled variable. Bold text indicates a substantive effect (CIs do not cross 0).

| | Estimate | 2.5% CI | 97.5% CI |
|---|---|---|---|
| **Intercept** | **1.89** | **1.57** | **2.22** |
| **Time to breed** | **-0.06** | **-0.10** | **-0.01** |
